# MSNovelist: *De novo* structure generation from mass spectra

**DOI:** 10.1101/2021.07.06.450875

**Authors:** Michael A. Stravs, Kai Dührkop, Sebastian Böcker, Nicola Zamboni

## Abstract

Structural elucidation of small molecules *de novo* from mass spectra is a longstanding, yet unsolved problem. Current methods rely on finding some similarity with spectra of known compounds deposited in spectral libraries, but do not solve the problem of predicting structures for novel or poorly represented compound classes. We present *MSNovelist* that combines fingerprint prediction with an encoder-decoder neural network to generate structures *de novo* from fragment spectra. In evaluation, *MSNovelist* correctly reproduced 61% of database annotations for a GNPS reference dataset. In a bryophyte MS^2^ dataset, our *de novo* structure prediction substantially outscored the best database candidate for seven features, and a potential novel natural product with a flavonoid core was identified. *MSNovelist* allows predicting structures solely from MS^2^ data, and is therefore ideally suited to complement library-based annotation in the case of poorly represented analyte classes and novel compounds.

## Introduction

A key challenge in mass spectrometry, particularly in metabolomics and non-targeted analysis, is feature annotation, or the assignment of chemical identity to unknown signals from their exact mass and fragment (MS^2^) spectra. Compounds may be identified by either searching against mass spectral libraries from synthetic standards, or by searching against data inferred from structures (so-called “*in silico* methods”). Both approaches have limitations. Matching against experimental libraries^1,2^ is limited by the actual availability of standards and curated spectral data which poorly represents the diversity and complexity of the chemical space. Searching against structural databases (e.g. PubChem^3^ or KEGG^4^) include non-trivial simulation of spectra^5^ or fragmentation patterns^6^ for candidate compounds, or the prediction of high-dimensional molecular descriptors for spectra^7–9^.

Importantly, none of these methods is able to identify truly novel and unexpected compounds like unknown natural products, drug metabolites, or environmental transformation products.

In principle, the simplest and entirely database-independent approach to assign structural identity to truly unknown compounds is to first determine the molecular formula, then enumerate all possible candidates, and finally score against experimental data^10–12^. This approach fails in practice because of the combinatorial explosion in the number of structures that can exist even for simple formulas^13^. Recent strategies for identification of true unknowns have instead relied on expanding compound databases using chemical reaction rules^14,15^, identifying partial structures using spectral networking^16^ and “hybrid search” (MS^2^ library search including mass shifts)^17^ or, recently, assigning chemical classes *in silico* using machine learning^18^.

In the context of computational drug design, deep learning algorithms for targeted *de novo* molecule generation have recently emerged. These methods allow querying the full chemical space without enumerating candidates. In analogy to the methods used for text generation, Gómez-Bombarelli et al. ^19^ used a variational autoencoder (VAE) with a recurrent neural network (RNN) to generate textual representations of molecules, i.e. SMILES^20^. Similarly, Segler et al. ^21^ used a RNN sequence model to generate molecules. Numerous variations of these models generate molecules in the form of SMILES, SMILES-related representations, or directly as graphs, achieving specific chemical properties by fine-tuning, optimization in latent space, or reinforcement learning, see e.g.^22–24^.

For mass spectrometry, two recent approaches have used molecule generation to generate candidate libraries based on a target collision cross-section (CCS) and mass^25^, or for a specific compound class^26^. However, these methods do not take the structural information from MS^2^ spectra in account, and therefore only provide a list of candidates that needs further filtering. If information from MS^2^ spectra could be used *directly* for *targeted* structure generation, this would bypass the combinatorial bottleneck for *de novo* structure elucidation. Unfortunately, using MS^2^ spectra to directly train molecule generation models is currently not feasible, because of the limited amount of training data (up to 30’000 molecules ^18^) which is more than an order of magnitude below standard requirements for generative models.

We tackle this fundamental problem by utilizing CSI:FingerID^8,9^ for molecular fingerprint prediction. CSI:FingerID predicts a high-dimensional molecular fingerprint from a query MS^2^ spectrum, then uses this fingerprint to query a molecular structure database. Such fingerprints can be deterministically computed for any molecular structure and independently from spectral libraries; hence, molecular fingerprints can be used to train any downstream method with virtually unlimited training data^18^. If the structural information in predicted fingerprints is sufficiently descriptive, it allows training a model to generate structures compatible with a specific MS^2^ spectrum, without requiring millions of labeled MS^2^ spectra as training data and without the need to constrain the search to a structural database.

Here, we present *MSNovelist*, a method for *de novo*, database-free structure elucidation that couples prediction of fingerprints from MS^2^ data with generation of structures from fingerprints. Our method uses the full probabilistic fingerprint obtained from CSI:FingerID to predict the probability of corresponding structural features directly from data. This allows us to exploit 1) probabilistic input and 2) fingerprint information which does not map directly to actionable constraints for structure generation. We evaluate the method’s performance in context of current state-of-the-art database search on a reference dataset from the GNPS spectral library, and on the CASMI 2016 structure identification challenge. As an exemplary application of de novo spectral annotation, we apply our method to a bryophyte LC-MS dataset and putatively annotate seven novel chemical structures.

## Results

### Overview of the approach

*MSNovelist* performs *de novo* structure elucidation from MS^2^ spectra in two steps (Figure 1).First, SIRIUS and CSI:FingerID are used to predict a molecular formula *x*^*M*^ and a structural fingerprint *x*^*F*^, respectively, from an MS^2^ spectrum as previously described^9^. Second, *x*^*M*^ and *x*^*F*^ are used as inputs for an encoder-decoder neural network, which predicts SMILES sequences based on the structural features described by the input fingerprint (Figure 2). Conceptually, the model learns how to represent the structural fingerprint features in a SMILES string.

**Figure 1:**
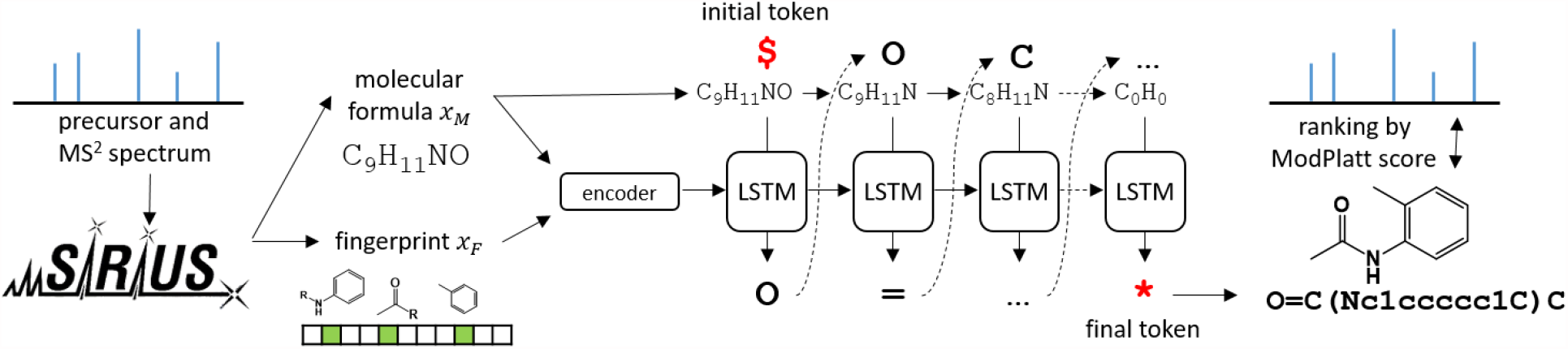
Conceptual overview of MSNovelist. Using the existing SIRIUS and CSI:FingerID approach, a molecular fingerprint and a molecular formula are predicted. This data is used as input to an encoder-decoder LSTM architecture to predict a SMILES sequence.

**Figure 2:**
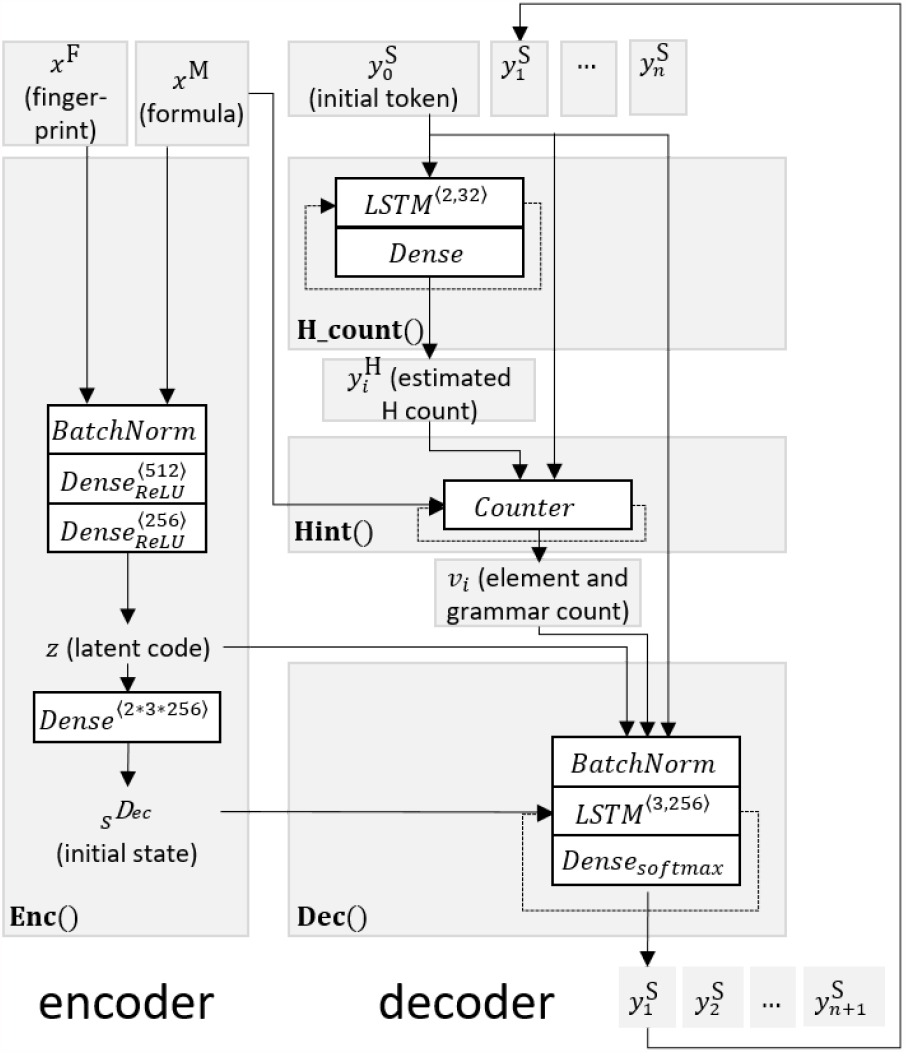
Model architecture. The encoder generates a reduced-dimensionality representation z from fingerprint x^M^ and molecular formula x^F^, and computes initial states for the decoder LSTM network. The decoder is composed from a recurrent neural network (Dec) which predicts an output token per timestep from the preceding token, the latent representation z, and an auxiliary vector v that counts remaining elements and open brackets in the SMILES code (Hint). An auxiliary LSTM network (H_Count) estimates hydrogen atom counts per token for use in the element counter. Concatenations are omitted in the scheme.

The encoder consists of three hidden layers and yields a real-valued vector *z*, which we consider the latent representation of the molecule. Vector *z* is further transformed via a single layer to starting state vectors *s*^*Dec*^ for the decoder LSTM. The decoder is a three-layer short-term-memory (LSTM) network^27^, which for any position *i* in a SMILES sequence, predicts probabilities for SMILES character *y*^*S*^_*i*_ and state 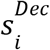 from an input of the previous character *y*^*S*^_*i*−1_, the preceding state 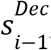, and the context vector *z*. This basic decoder model can be extended with additional information from an augmented feature vector *v*_*i*_ to increase performance. Vector *v*_*i*_ contains the running count of remaining atoms per chemical element, i.e. the atom count in *x*^*M*^ minus the sum of atoms of this element in the partial sequence up to *y*_*i*′_ and the number of open brackets, i.e. the count of open minus closing brackets in the partial sequence. This auxiliary vector aids the generation of syntactically correct SMILES, and molecules of a particular formula, since sequence termination is contingent on |*v*_*i*_| = 0 in the training set. Vector *v*_*i*_ is directly given from the SMILES sequence for heavy atoms, whereas the number of (implicit) hydrogen atoms is not directly evident in a partial SMILES. To this end, an auxiliary two-layer LSTM predicts a sequence of hydrogen atom counts per SMILES character, trained such that their sum matches the total hydrogen count in the molecule.

For sequence prediction, *x*^*F*^, *x*^*M*^ are encoded into *z* and 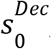, and the top-*k* sequences are decoded via beam search. For every query tuple (*x*^*F*^, *x*^*M*^), the model returns *k*′ ≤ *k* valid structures *S*^(1..*k*′)^. To find the structure with the best match to the query fingerprint, the structures were eventually ordered by the modified Platt (ModPlatt) score^8^, which measures the match of a predicted fingerprint to a deterministically computed fingerprint of a candidate structure. Further details are provided in the Online Methods.

### Method validation

The encoder-decoder model was trained on a dataset of 1 232 184 chemical structures compiled from the databases HMDB 4.0^28^, COCONUT^29^ and DSSTox^30^, and 14 047 predicted fingerprints for predicted fingerprint simulation (see SI for details). Importantly, all structures used for fingerprint simulation or present in evaluation datasets (see below) were removed from the training set to effectively evaluate the ability to identify structures that have not been observed before. The full structural prediction model was validated using 3863 MS^2^ spectra from GNPS^31^ closely matching the evaluation setup by Dührkop et al.^8,9^ These spectra cover heterogeneous types of compounds, samples, instrumentation, spectral quality, etc. No additional quality control was performed before subjecting these spectra to *de novo* annotation. For every spectrum, we extracted the top-128 SMILES sequences for fingerprints predicted using structure-disjoint tenfold cross-validation. As the RNN can generate invalid SMILES, or also generate different SMILES strings that encode for the same structure, we verified and dereplicated molecular structures. At least one valid structure was generated for all but one MS^2^ spectra. A structure with the correct molecular formula was generated for 99.5% of instances. The correct structure was retrieved for 45% and ranked first for 25% of the instances (Fig. 3a). In comparison, database search with CSI:FingerID was able to rank the correct structure on top for 39% of the spectra. In this subset of spectra which were correctly identified by CSI:FingerID (GNPS-OK, 1507 spectra; Supplementary Table 2), *MSNovelist* correctly retrieved 68% of the true structures, and in 61% (out of 1507) as the top candidate (Fig. 3b). In the cases where the true structure was not ranked first by *MSNovelist*, the generated structures were frequently very similar to the target molecule. This is shown by ten randomly chosen examples in Fig. 3e and Supplementary Fig. 1. Seven mispredictions were close isomers of the correct structure, one instance showed a partial mismatch in the skeleton, and only two predictions were completely wrong.

**Figure 3:**
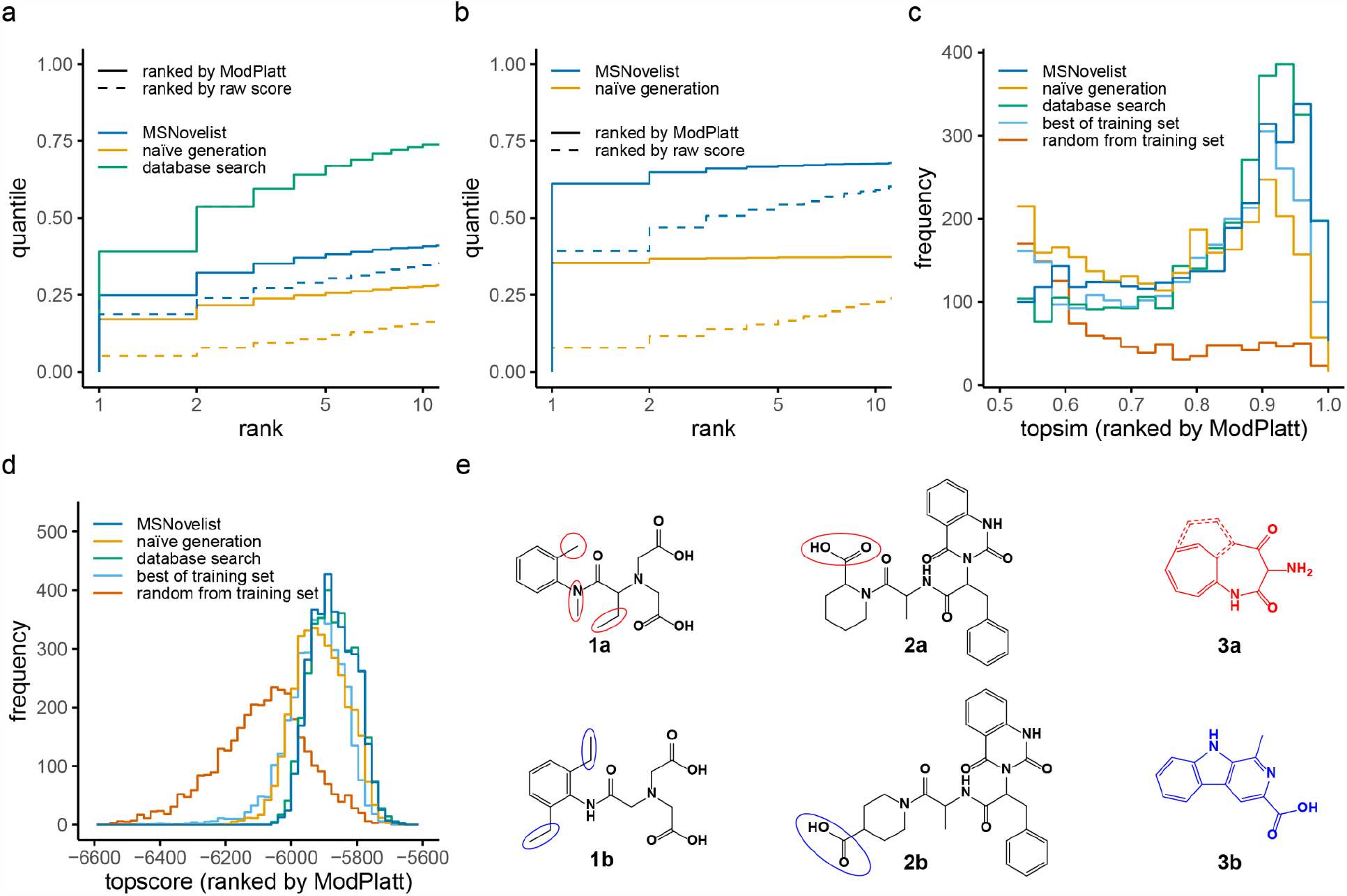
Validation of MSNovelist with GNPS dataset. a) Rank of correct structure in results for MSNovelist (blue), and naïve generation (orange), with ranking by ModPlatt score (solid) or ordered by model probability (dashed), and comparison to database search (CSI:FingerID on PubChem; green) for the GNPS dataset (n=3863). b) Rank of correct structure in results for MSNovelist and naïve generation, with ranking by ModPlatt score or ordered by model probability, and comparison to database search for GNPS-OK dataset (n=1507). c) Tanimoto similarity of best incorrect candidate to correct structure (topsim) for MSNovelist, naïve generation, database search, best candidate from training set, and random candidate from training set. d) ModPlatt score of top candidate (topscore) after reranking, for MSNovelist, naïve generation, database search, best candidate from training set (light blue), and random candidate from training set (red) e) Three randomly chosen examples of incorrect predictions (top candidate) from GNPS dataset. Structures 1,2,3a: de novo prediction; structures 1,2,3b: correct result. Red color marks sites predicted incorrectly by the model (or the entire molecule if the prediction was completely wrong), blue color marks the corresponding correct alternative.

The importance of structural information was tested with a model that lacked the fingerprint input to the encoder. This naïve generator creates structures with a specific molecular formula, but cannot make use of structural information. Importantly, the results of the naïve generator were again reranked by the ModPlatt score. In the GNPS dataset, naïve generation retrieved only 31% of all correct structures, and 17% were ranked first. While this is clearly lower than guided *de novo* generation, it also shows that deceptively high performance can be achieved purely by sampling large numbers of molecules and *a posteriori* reranking using the ModPlatt score. This highlights the need to consider such baselines for evaluation. A similar pattern is observed for the GNPS-OK subset, with 37% structures retrieved, 35% of which were ranked at the top. In both datasets, inclusion of structural information from fingerprints in *de novo* generation increased the recovery of true structures by 13 to 31 percentage points.

As a further benchmark, we attempted to generate structures for the 127 MS^2^ spectra from the CASMI 2016 competition^32^, which is a common benchmark for structure elucidation (Supplementary Fig. 2, Supplementary Tables 3 and 4). *MSNovelist* retrieved 57% of structures, with 26% ranked first. The naïve model achieved 52% retrieval, and 24% top-1 hits. From the 47 instances that were correctly identified by database search (CASMI-OK), *MSNovelist* retrieved 74% and identified 64% as first rank; exhibiting a wider gap to naïve generation (56% retrieved, 51% top 1). The improvement over the structurally naïve generator is marginal, indicating that the molecules in the CASMI dataset are similar to the training set and *a priori* likely under the model. Overall, these tests demonstrate that *de novo* structural annotation is in principle possible and in 50-70% of the cases generates candidates that are consistent with true structure.

### Extrapolation performance, chemical space coverage and model score

We conducted further evaluations to demonstrate model performance and to verify that *de novo* structures outscore training set structures in terms of similarity to the query fingerprint and molecule. First, we compared the chemical similarity of the best incorrect prediction to the correct structure (Fig. 3c). This allows a direct comparison of *de novo* predictions with the training set data, which contains only incorrect results (compare also Cooper et al.^33^). The best incorrect *de novo* prediction scored higher than best-in-training-set and nearly as high as the best incorrect database structure. This demonstrates that the model systematically generates combinations of chemical features not seen in the training set. In contrast, naïve generation slightly underperformed the best training set candidate, as expected from generation without structural guidance.

The ModPlatt score quantifies the match of the generated structure to the input fingerprint. Therefore, it directly measures the ability of the model to generate structures with desired chemical features independently of errors in fingerprint prediction. For the best *MSNovelist* candidates, the ModPlatt scores were essentially identical to the best database compounds and higher than best compounds in the training set (Fig. 3d). Again, the naïve model performed slightly worse than the best training set candidate. Finally, chemical space coverage was assessed by examining the rediscovery of high-scoring database compounds. Our *de novo* generation reached a mean of 58%, compared to 31% for naïve generation. We also evaluated the relevance of posterior reranking with the ModPlatt score. The raw model without reranking reached notable 19% and 39% correct (top-1) identifications in the GNPS and GNPS-OK datasets, respectively (Fig. 3a, 3b). The best compound identified by the raw model outscored the best training set compounds in the topscore benchmark, and is on par in terms of chemical similarity between the best incorrect prediction and the correct structure (Supplementary Fig. 5). This indicates that the raw score obtained by the model is already informative about structure-spectrum match. However, the use of the ModPlatt score for final ranking yields more correct identifications, and higher scores in the auxiliary benchmarks.

Similarly, we examined the necessity of element counting and hydrogen estimation for generation of valid results and their effect on model performance (Supplementary Fig. 3, 4; Supplementary Tables 1-4). In summary, the model was still able to produce high-scoring results without the additional components, they increased the number of valid results with the correct molecular formula, and consequently, slightly improved overall retrieval and chemical space coverage.

Finally, we examined the impact of the number of generated candidates. Generation of only 16 candidates by *MSNovelist* was sufficient to outcompete top-128 naïve candidates in all metrics for all four datasets and subdatasets (see Supplementary Fig. 2, 5 and Tables as noted above). This provides further evidence that *MSNovelist* directly generates structures with high spectrum-structure match without requiring extensive sampling.

### De novo annotation of bryophyte metabolites

An objective of *de novo* annotation in discovery metabolomics is to identify novel biological small molecules. We demonstrate this application for a dataset of nine bryophyte species (Peters et al.^34^) Bryophytes (“mosses”) are known to produce diverse secondary metabolites, but are not extensively studied, presenting a likely opportunity for natural product discovery. From the data repository (MTBLS709), we extracted 576 consolidated MS^2^ spectra and with SIRIUS predicted molecular formulas and fingerprints, and obtained structure candidates from comparison with a database search. For spectra where the molecular formula could be predicted with high confidence (≥ 80% explained peaks, ≥ 90% explained intensity, ≥ 0. 9 ZODIAC score; 224 spectra), we predicted structures *de novo* with our method and compared ModPlatt scores for the best *de novo* and database candidates. (Fig. 4a). Points above the diagonal indicate that the *de novo* structure is a better fit to the spectrum than any database entry. In 27 cases the same structure was identified with both approaches. For 169 cases (75%), the *de novo* structure scored higher than the database. Only for 55 cases the database structure scored higher.

**Figure 4:**
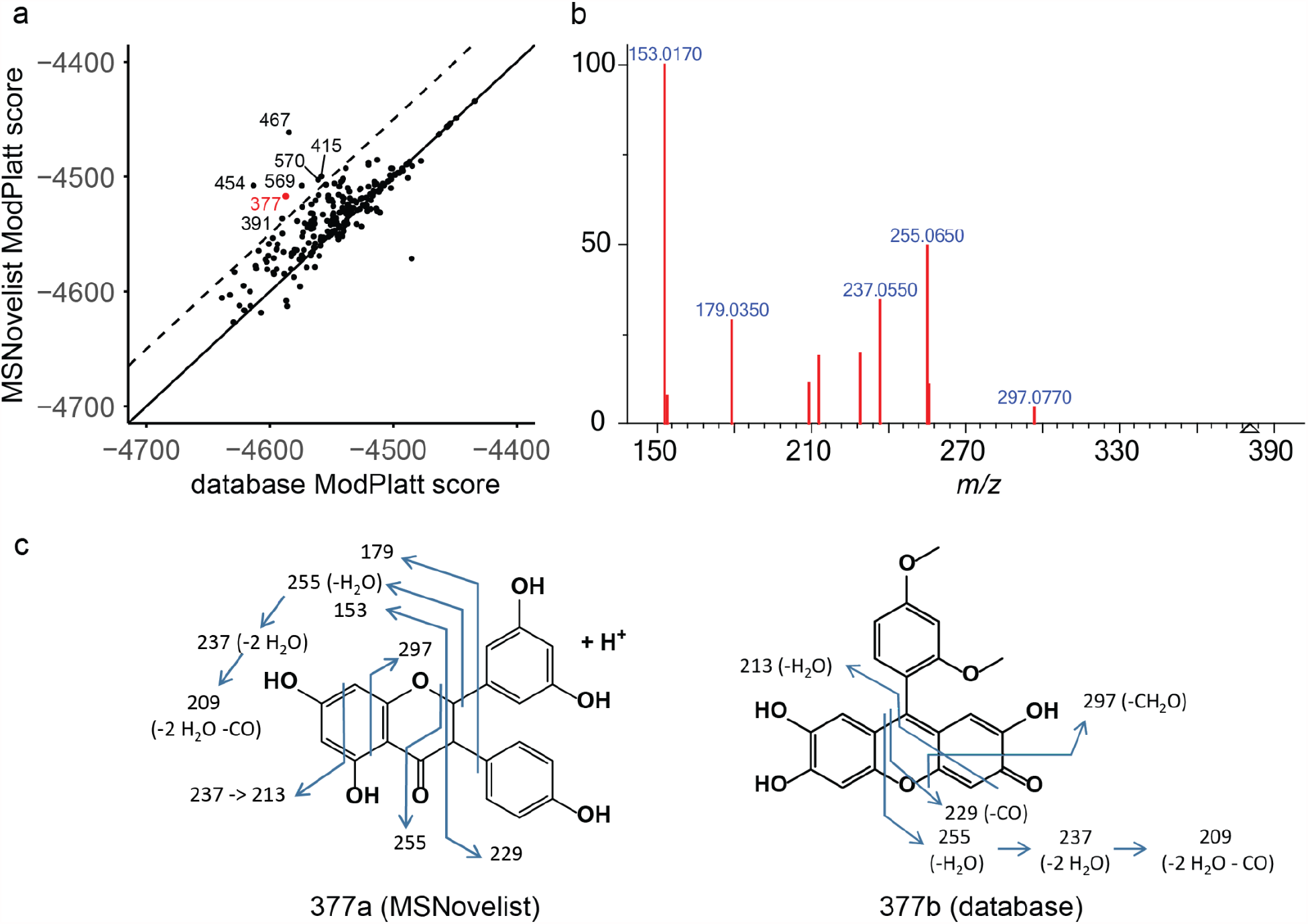
De novo annotation of bryophyte metabolites. a) Scores of best MSNovelist candidates versus best database scores for 232 spectra; regular line: 1:1 line; dashed line: ModPlatt_MSNovelist_ = ModPlatt_DB_ + 50; labels: spectrum ID. b) MS^2^ spectrum of feature 377. c) Proposed spectrum interpretation for structure 377a (MSNovelist) and 377b (database).

We inspected in detail the 7 cases with prominent differences between *de novo* and database score (Supplementary Table 5). Details are presented in Supplementary Tables 5-13 and Supplementary Fig. 6. For the example of feature 377 (*m/z* 381.1020, C_21_H_16_O_7_), the *de novo* predicted structure 377a is a polyphenolic structure with a flavonoid core (Fig. 4c) and all observed fragments (Fig. 4b) are consistent with the proposed structure. Fragment 153 and neutral loss 126 (fragment 255) are shared with the flavonoid hesperetin. Seven peaks are shared with the structurally similar chrysin-7O-glucuronide and all matching peaks relate to the aglycon. In contrast, the best database candidate fails to explain peaks 153 and 179. The structure predicted *de novo* is similar to known natural products formosumone A and struthiolanone. The biosynthetic origin of a C_21_ polyphenol remains unclear, but we hypothesize that it could arise from a condensation as in the case of struthiolanone^35^. We noted that multiple alternative *de novo* structures also strongly outscored the best database suggestion and are compatible with the observed spectrum. In any case, there is solid evidence that feature 377 is a novel natural product with a flavonoid core and an interesting candidate for further structure and biosynthetic pathway elucidation.

In summary, for four of the seven cases the *de novo* suggestions are better at explaining the MS^2^ spectrum, and one is equally good, proposing multiple avenues for natural product discovery in a relatively unstudied phylum. The proposed structures should be seen as starting points for further investigation, e.g. by targeted acquisition and analysis of more fine-grained MS^2^ data including higher-mass fragments, but importantly also by isolation and characterization by NMR, and elucidation of their biosynthesis.

## Discussion

*MSNovelist* demonstrates that *de novo* annotation of MS^2^ spectra is possible, challenging the existing paradigm that the complexity of small-molecule chemical space precludes these approaches. *MSNovelist* constitutes the first direct application of a chemical generative model to mass spectrometry data. While deep learning models have previously been used to generate candidate libraries to use with independent methods for structure identification by MS^2 25,26^, *MSNovelist* is capable of integrating the structural information encoded in probabilistic fingerprints. Given that certain isomers fragment (almost) indistinguishably, structural elucidation from this data is clearly not possible in all cases; yet, *MSNovelist* suggested reasonable molecular structures for more than half of the MS^2^ spectra.

Three aspects made this achievement possible. First, structural fingerprint predictions from MS^2^ spectra (by CSI:FingerID) directly encode structural information and may act as a blueprint for a molecule. Second, we decoupled MS^2^ interpretation from structure generation, allowing us to train the generative model with millions of structures^18^ and independently from experimental MS^2^ data. Third, we exploited the analogy between writing a SMILES code based on a chemical fingerprint and image captioning, i.e. writing a descriptive sentence based on a feature vector. In this context, *de novo* structure elucidation is interpreted as a translation-like task from fingerprint to structure. Trained in this manner, the model works independently of pre-existing scores for spectrum-to-structure matching; though we acknowledge that final reranking with the ModPlatt score is important for best results.

*De novo* annotation could alternatively be treated directly as an optimization task with a pre-existing scoring function; i.e., candidate structures can be obtained by traversing the latent space in a generative model or with reinforcement learning on the score. This would not require input directly informative about structure (such as fingerprints), and would therefore work with any spectrum-structure score, such as match to simulated MS^2^ spectra using CFM-ID^5^ or any other score used by database search approaches. It would also allow the integration of orthogonal information, most trivially retention time prediction. However, optimization-based approaches rely on external, expensive oracles (e.g., the fingerprinting implementation required for ModPlatt computation is slow compared to sequence generation). We experimented with reinforcement learning based on the REINVENT model^23^. While the approach is promising, the integration of fingerprint fidelity and molecular formula match objectives, as well as other scores, can lead to undesired tradeoffs and will require further research.

From the point of view of chemical generative models, the task to generate structures compatible with a fingerprint resembles Tanimoto and Rediscovery benchmarks (e.g. in the Guacamol benchmark suite^24^) with the added complications that we are working with fuzzy, incomplete fingerprints, and always require precise isomers of a single molecular formula. Given existing results with these tasks, it is not surprising that the model is able to generate structures with high spectra-structure match scores. It was, however, unclear whether the fuzzy predicted fingerprint input would provide constraints narrow enough to enable structure elucidation. We showed that our model retrieves a large proportion of correct hits, as well as additional incorrect structures that score highly in database search. Further, incorrect high-scoring structures were highly similar to the correct answers, both anecdotally and by evaluation. Given how vast we usually perceive small molecule chemical space to be^36^, the results for chemical space coverage and ranking appear better than naïvely expected. Further, a notable part of results could be rediscovered even by isomer sampling, without structural fingerprint input. This indicates that the chemical space described with the present model and training set is comparably well-confined. This might limit the model’s ability to discover chemistry extremely different from known molecules; however, such discoveries would be of limited use in practice, since a practitioner would be unlikely to trust such structure suggestions. Even with this “conservative” model of chemical space, we were able to predict plausible novel molecules in biological datasets. Finally, we note that the SMILES-based model used in the present work is considerably simpler than state-of-the-art models, which generate graphs directly (e.g., ^37–40^), and we did not apply robust alternatives to SMILES such as DeepSMILES^41^ and SELFIES^42^. While more advanced models may be applied for future approaches, the choice of chemical model is not central to the presented work. We could show that even with such a “simple” model, good results can be obtained for a practical task; and we consider our work also a baseline for more elaborate developments.

In conclusion, this work contributes a further building block to the growing set of methods for untargeted computational mass spectrometry. MS^2^ library search and database search will remain workhorses for high-throughput annotation, and systems-oriented methods^18,43,44^ are becoming important for a view centered on broader categories rather than individual structures. In contrast, *de novo* methods might become relevant for discovery applications and structure hypothesis generation. We foresee that integrating prior knowledge (e.g., information about known parent compounds in a metabolism study, or about known biological and chemical composition of a sample) will help to increase the accuracy and practical utility of *de novo* annotation.

## Acknowledgements

The authors thank Marcus Fleischauer for support with the CSI:FingerID webservice and SIRIUS, and the HPC team of the ETH Zurich Scientific IT service for providing cluster support.

## Code and data availability

*MSNovelist* is available on Github, https://github.com/meowcat/MSNovelist. The dataset MTBLS709 analysed during the current study is available in the MetaboLights repository, https://www.ebi.ac.uk/metabolights/MTBLS709. Processed data is available on the GNPS repository, https://gnps.ucsd.edu/ProteoSAFe/status.jsp?task=b8b481147b844ebda2481bf9656baec8. The HMDB and COCONUT databases are available on Zenodo, https://zenodo.org/record/3375500 and https://zenodo.org/record/3778405. The DSSTox database is available at ftp://newftp.epa.gov/COMPTOX/Sustainable_Chemistry_Data/Chemistry_Dashboard/MetFrag_metadata_files/CompTox_17March2019_SelectMetaData.csv. All further data is available from the authors on reasonable request.

## Author contributions

MS conceived the idea, implemented, trained and evaluated the model. KD and SB developed the fingerprint simulation, processed the training data, and contributed to the evaluation method. NZ supervised the study. All authors contributed to writing the manuscript.

## Online Methods

### General

For all purposes, we consider chemical structures unique, ignoring stereochemistry, when they have an identical InChIKey2D (see below). For any unique structure, one or more SMILES codes (of stereoisomers, or also multiple spectra for one isomer) may exist in our datasets. Therefore, datasets were split into folds considering unique structures, such that no unique structure was present in test and training sets; i.e., they were “structure-disjoint”.

We refer to deterministically calculated structural fingerprints for a molecule as *structural fingerprints* (struct-FP), and to fingerprints predicted from an MS^2^ spectrum with CSI:FingerID as *spectrum fingerprints* (spec-FP). Fingerprints that were predicted with CSI:FingerID in a 10-fold structure-disjoint cross-validation setup, so that the structures of all instances are unknown to CSI:FingerID, are called *cross-validated spectrum fingerprints* (CV-spec-FP). Fingerprints generated by perturbation from struct-FP to simulate spec-FP are called *simulated fingerprints* (sim-FP).

### Definitions

We denote *Dense*^<*n*>^ a dense (fully connected) layer with *n* units (and linear activation). We denote 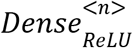 a corresponding layer with the activation function *ReLU*(*x*) = *max*(0, *x*). We denote *LSTM*^<*n,m*>^ a LSTM recurrent neural network with *n* layers and *m* units per layer as described by Hochreiter and Schmidhuber^27^ and implemented in Keras/Tensorflow, with *tanh* activation, sigmoid recurrent activation, and no dropout.

We denote *Counter*_*M*_ the (recurrent) countdown function

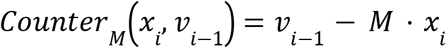

with a starting state *v*_0_ ∈ *R*^*n*^ and a (constant, non-trainable) matrix *M* ∈ *R*^*n,dim*(*x*)^. Typically *M* ∈ {1, 0, − 1}^*n,dim*(*x*)^.

### Dataset and data preprocessing

A training set was composed from the databases HMDB 4.0^28^, COCONUT^29^ and DSSTox^30^. The training set was filtered to remove molecules which couldn’t be parsed with RDKit, SMILES codes longer than 127 characters, disconnected SMILES codes (containing a dot), molecular weight larger than 1000 Da, a formal charge, more than 7 rings (as specified in SMILES), or elements other than C, H, N, O, P, S, Br, Cl, I and F. All structures contained in the CV-spec-FP dataset (see below) were removed from the training set. Finally, the training set contained 1 232 184 molecules with 1 048 512 distinct structures (by InChIKey2D), and was split into ten structure-disjoint folds.

For generation of sim-FP (see below) and model evaluation, a dataset of 14047 CV-spec-FP was obtained, corresponding to the openly available part of the CANOPUS^18^ evaluation data, the GNPS dataset, and the CASMI dataset. The CV-spec-FP dataset was split into ten structure-disjoint folds, and (arbitrarily) matched to one fold in the training set. All structures present in the CASMI dataset were assigned to the same fold, such that the dataset is completely unknown to the corresponding model.

All molecules in input data were initially retrieved as SMILES code or InChI code. For every molecule, a SMILES string standardized with the PubChem standardization service was retrieved. Using RDKit, the structure was parsed, the InChIKey was generated, and the first 14 characters (a hash describing atom connectivity ignoring stereochemistry and charge, “InChIKey2D”) was extracted. For every unique InChIKey2D, a Pubchem-standardized SMILES string^45^ was retrieved, from which stereochemical information was removed using regular expressions. The resulting stereochemistry-free SMILES was processed in Java using the CDK toolkit (version 2.3) and SIRIUS libraries (version 1.4.3-SNAPSHOT at the time of writing) to obtain an aromatic canonical SMILES code and a >8000-bit struct-FP as described elsewhere containing CDK Substructure fingerprints, PubChem fingerprints, Klekota-Roth fingerprints^46^, FP3 fingerprints, MACCS fingerprints, ECFP6 topological fingerprints^47^ and custom rules for larger substructures^18^. The aromatic canonical SMILES code was parsed with the toolkit RDKit, and the molecular formula extracted. The struct-FP, SMILES code and molecular formula were stored in a database, or in a CSV-formatted text file. For the CV-spec-FP dataset, the CV-spec-FP was additionally stored. The multikernel SVM method employed by SIRIUS predicts (at the time of writing) 3609 bits from the >8000-bit struct-FP; in the following, the struct-FP shall denote only these 3609 bits, and *FP*(*S*) shall denote the function that calculates the 3609-bit struct-FP for a chemical structure *S*.

### Input processing and encoding

For training, sim-FP were generated on the fly (during training) from struct-FP by random sampling from the CV-spec-FP dataset (minus the current training fold), using a procedure similar to the description in Dührkop et al.^18^ (“first method”), as shown in Algorithm 1.

#### Algorithm 1

Fingerprint simulation

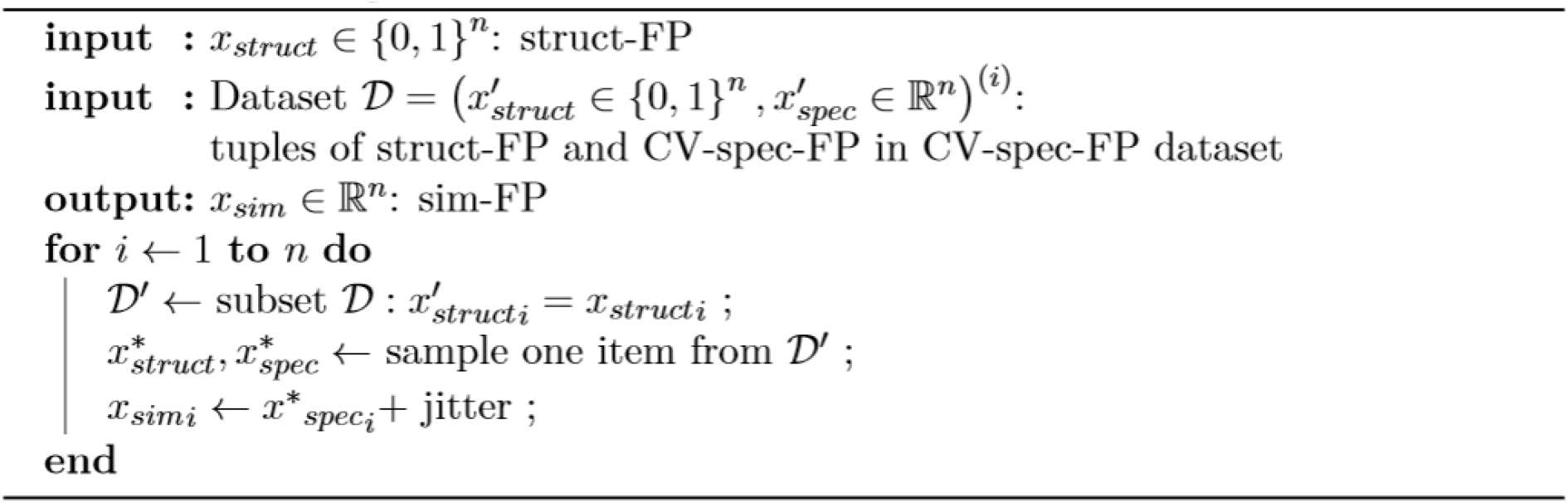

The sim-FP were used either verbatim (probabilistic input) or rounded to 0 or 1 (discrete input). Discrete input was chosen in the final model because it led to superior results in chemical space coverage and correct structure retrieval. For evaluation and in inference mode, spec-FP were also correspondingly rounded.

Additionally, a method based on correlated sampling (similar to “second method”, Dührkop et al.^18^) was implemented, which takes into account correlations between fingerprint bits. However, the method led to identical results in chemical space coverage and correct structure retrieval; therefore, the simpler method was further used.

The molecular formula was encoded as a vector *x*^*M*^ ∈ *N*^*m*^ for the *m* = 10 elements *E* ∈ *C, F, I, Cl, N, O, P, Br, S, H* (with *x*^*M*^_*i*_ denoting the sum of atoms of element *E*_*I*_ in the molecule). We denote *MF*(*S*) the function returning the molecular formula for a structure *S*.

The aromatic canonical SMILES codes were split into tokens consisting of a single character (e.g. `C,c,=,N,3`), the two-letter elements `Br,Cl` substituted as `R,L`, or a sequence of characters delimited by square brackets (e.g. `[nH],[N+]`), denoting special environments. *t* = 36 tokens occurring >100 times in the training set were retained, the remaining tokens (20 tokens with a total of <2000 occurrences) were discarded and ignored. All token sequences were prefixed with a start token (`$`), postfixed with a final token (`*`) and padded to a fixed length of *l* = 128 with a pad token (`&`). The sequences *s* were then transformed to a one-hot encoded matrix *Y*^*S*^ ∈ {0, 1}^(*l,t*)^ (such that *Y*^*S*^ _*i,j*_= 1 ⇔ *s*_*i*_ = *j*) with column vectors *y*^*S*^_*i*_.

### Data augmentation: Element count and grammar balance

For promoting the formation of correct SMILES and the correct chemical formula, the input vector was augmented with a counter *v*_*i*_. This vector counted the remaining atoms per element, and open parentheses in the current sequence, starting from the molecular formula *x*^*M*^ (and zero open brackets) as the initial state. Formally,

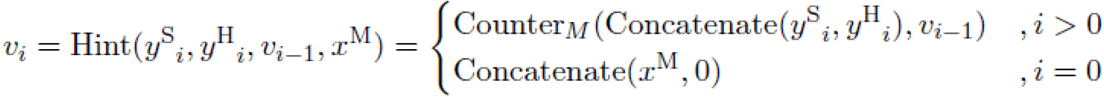

The counter matrix *M* consisted of an upper part mapping input tokens, a row mapping the predicted implicit hydrogen count to the hydrogen element, and a row mapping tokens (,) to 1, − 1 respectively:

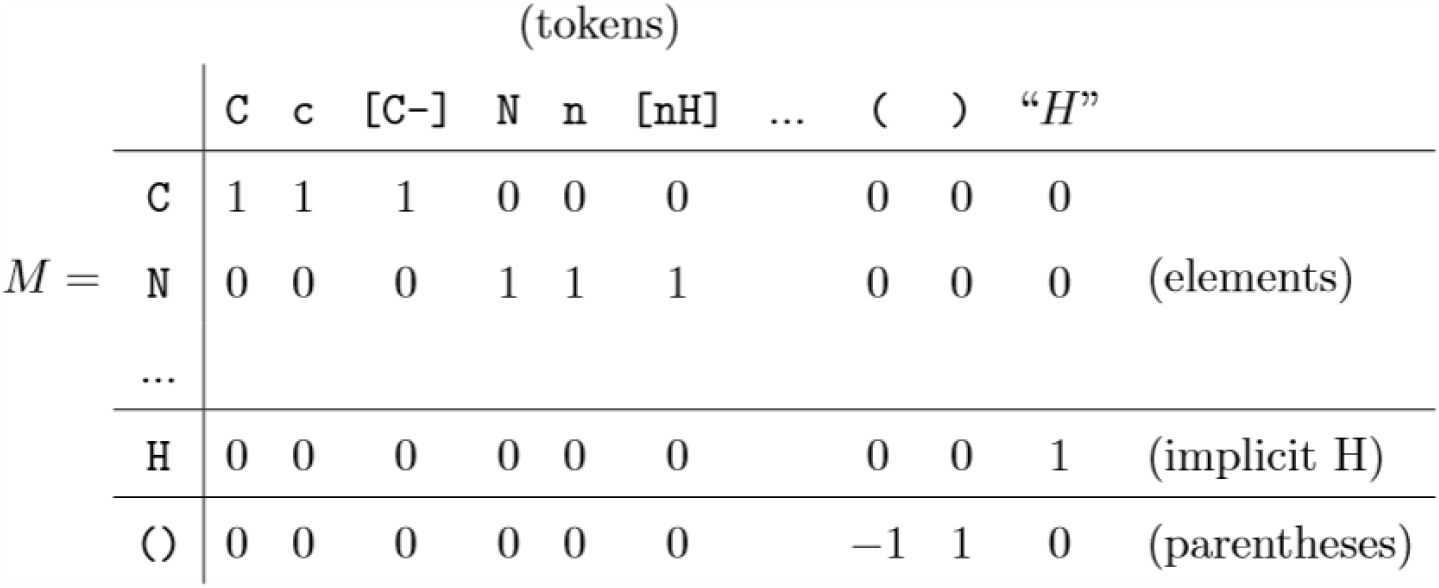

### Data augmentation: Hydrogen count estimation

As opposed to heavy atom counts, which are directly specified by tokens in the SMILES sequence, hydrogens are implicitly described, and assigned after constructing the molecular graph. For use in data augmentation (see above), we estimated implicit hydrogens for every token in a partial SMILES sequence from the sequence context. We trained an LSTM network with parameters ϕ

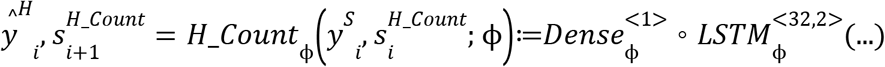

where the output *y*^*H*^ _*i*_ is the estimated hydrogen count for token *i*. Instead of deriving the hydrogen count per sequence element from actual molecular graphs for training, we summed 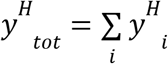 and minimized the loss 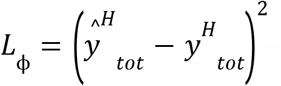, such that the sum of hydrogens assigned to each sequence element (ignoring termination and padding tokens) matches the total count in the molecule given by *x*^*M*^. *H*_*Count* is trained concomitantly, but separately from the remaining network (no gradients propagate through *y*^*H*^). We note that since only left-hand context is available, “hydrogen equivalents” can be positive or negative numbers (e.g., a branch opening (may contribute a negative hydrogen.)

### Fingerprint encoder and sequence decoder

The encoder block *Enc*(*x*^*F*^, *x*^*M*^; θ) consists of a batch normalization layer and two dense layers (512 and 256 units, respectively; ReLU activation) to compute a latent code *z* from the concatenation of inputs *x*^*F*^ and *x*^*M*^, and a dense layer (2*3*256 units, linear activation) to compute 2*3 initial states 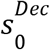 for 3 LSTM layers of 256 units each from the latent code *z*.

Similar to related work, the decoder was implemented as an LSTM with three layers of 256 units per layer and a final dense layer with the number of output tokens. The input to the LSTM consists of the context vector *z* (constant over the sequence), the preceding sequence token *y*^*S*^_*i*′_ the molecule target vector *v*_*i*_ and the LSTM state 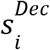:

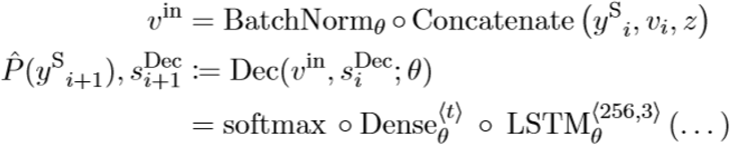

Parameters θ (for *Enc, Dec*) and ϕ (for *H*_*Count*) were found (in parallel, but independently) through training in teacher forcing manner^48^ to minimize the categorical crossentropy loss.

As in common image captioning models^49^, the latent space is not explicitly regularized; the translation task (from fingerprint features to SMILES representation) is expected to be a bona fide regularizer, given a small enough latent space. In variational inference, (over)regularized models with complex decoders (particularly when trained with teacher forcing) tend to ignore latent code^50,51^. This may be acceptable in tasks where a higher diversity of results is desired. However, the present task requires decoding to be as precise as possible. Multiple regularized models were additionally examined: for example a variational autoencoder (VAE)-like model regularized with Kullback-Leibler divergence (KL-VAE) or with the more information-preserving maximum mean divergence (MMD-VAE); and a model regularized by imposing the additional objective of reconstructing the true structural fingerprint of a compound from a spectrum-predicted fingerprint. All models with additional regularization performed worse than the base model.

### Implementation and training details

The model was implemented in Keras / Tensorflow, version 2.4.1 on Python 3.7. The network was trained with stochastic gradient descent using the Adam optimizer^52^ with a learning rate of 0.001, β_1_ = 0. 9, β_2_ = 0. 999 and ϵ = 10^−7^ over 30 epochs. Though the evaluation loss on sim-FP continued to minimally improve up to epoch 30, the evaluation performance of models did not improve or decrease meaningfully anymore after approximately 15 epochs. For evaluation, the weights after 20 epochs were used for all models. (Note: Since weights were only stored if loss had improved over the last epoch, not all folds have a weight at epoch 20; in this case, the last preceding weight was used.) The model was trained on a HPC cluster on a GPU node, using one dedicated Nvidia GTX 1080 or Nvidia GTX 1080 Ti GPU, 5 cores of a Xeon E5-2630v4 processor, and 80 GB RAM. Training time was approximately 45 min per epoch.

### Prediction

For sequence prediction from spec-FP, the latent code *z* and starting states 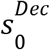 were predicted with the encoder. For hydrogen prediction, 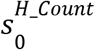 was initialized with zeros; for formula/grammar hinting, *v*_0_ = *Concatenate* (*x*^*M*^, 0) as stated above. Given *z* and the combined initial state *s*_0_ = (*s*^*Dec*^, *s*^*H_Count*^, *v*)_0_, a beam search with beam width typically *k* = 128 was performed with the decoder as described in Algorithm 2, with *argtop*_*k*_ (*x*) the positions of the top-*k* elements in vector *x*. In the implementation, the decoding is performed in parallel for multiple queries.

For evaluation, structure-disjoint cross-validation was used; i.e., the model used for structure prediction for any CV-spec-FP was trained without any fingerprints for this structure in the CV-spec-FP dataset used for fingerprint simulation.

For ablation studies, stochastic decoding was additionally used. Here, *k* sequences are sampled independently. Starting with the initial token, and the given (deterministic) starting state, a token *y*_*i*+1_ = *t* is sampled according to its probability distribution 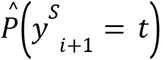, until the termination token is sampled.

### Modified Platt score

The ModPlatt score^8,53^ measures the match between a spec-FP *x*^*F*^ ∈ *R*^*n*^ and a struct-FP *y*^*F*^ for a structure *S*; *y*^*F*^ = *FP*(*S*) ∈ {0, 1}^*n*^, taking into account the predicted Platt probability (after additive smoothing) and the CSI:FingerID prediction statistics for each bit. The sensitivity *a*_*i*_ = *TP*_*i*_ /(*TP*_*i*_ + *FN*_*i*_) and specificity *b*_*i*_ = *TN*_*i*_ /(*TN*_*i*_ + *FP*_*i*_) (with *TN* the true negatives, *TP* the true positives, *FP* the false positives and *FN* the false negatives for bit *i*, respectively) are obtained from CSI:FingerID output. The ModPlatt score is then calculated as follows:

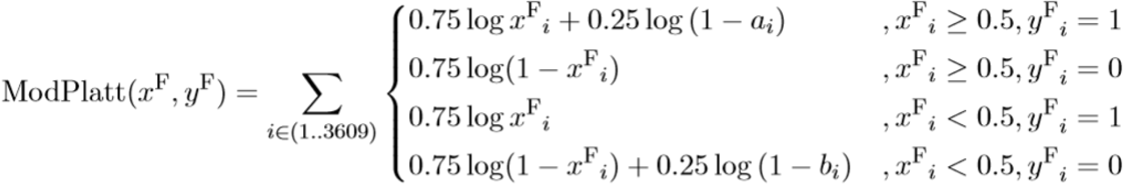

#### Algorithm 2

Beam search

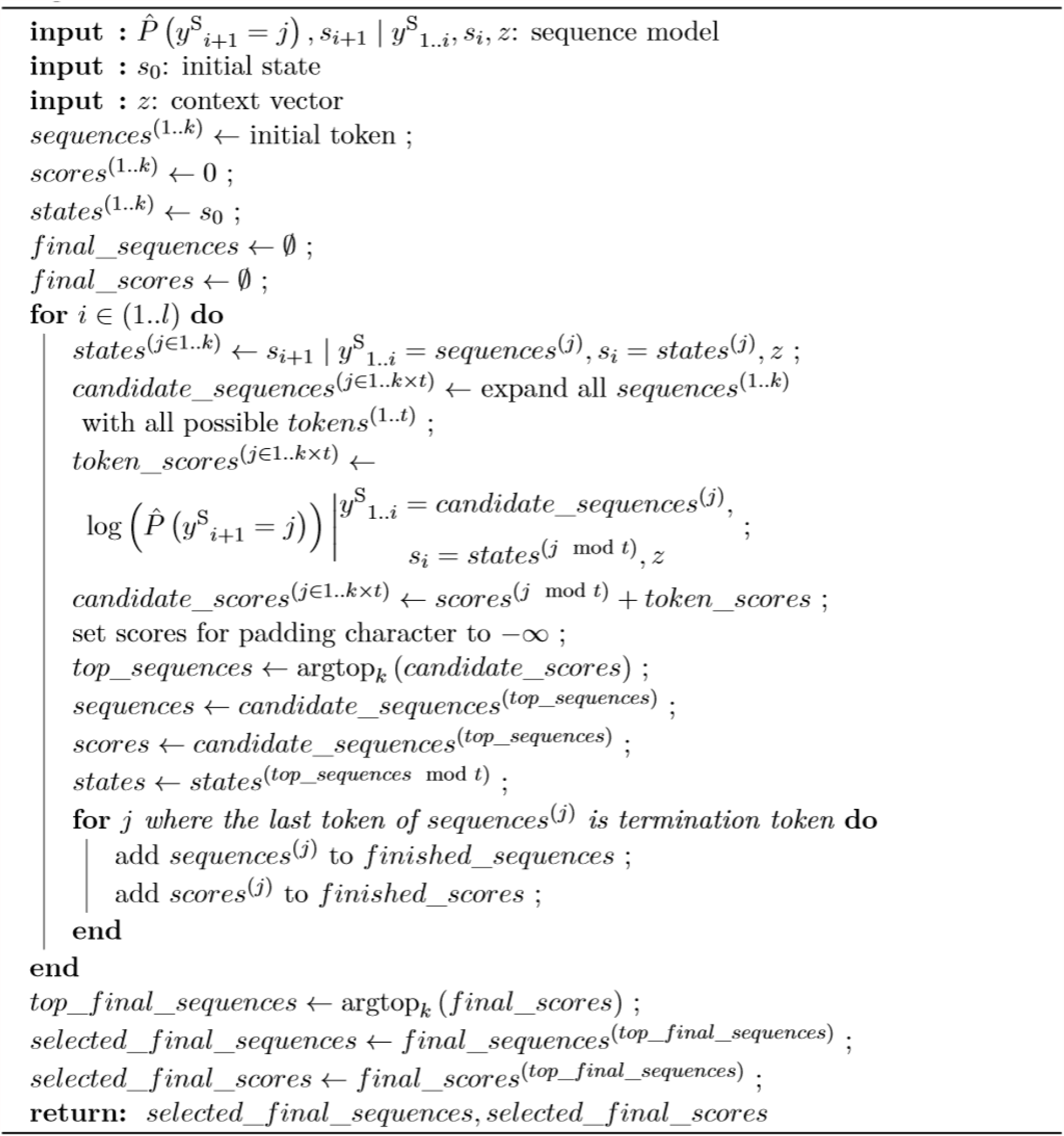

### Evaluation metrics

The model and the corresponding baselines were compared with multiple metrics. The following scores were calculated for every instance of a dataset, and their median and first and third quartiles, and/or their histograms, were reported.

- “% valid SMILES”: For every instance of a dataset, the percentage of predicted sequences that could be successfully parsed to a molecule using RDKit without any modifications.
- “% correct MF”: For every instance of a dataset, the percentage of predicted sequences that could be successfully parsed to a molecule using RDKit without any modifications, *and* that additionally matched the molecular formula of the correct structure.
- “topscore”: For every instance of a dataset, the ModPlatt score of the highest-ranked candidate versus the query fingerprint.
  ‐ ranked by raw score: the ModPlatt score of the highest-ranked candidate according to the model score, i.e. the probability under the model
  ‐ ranked by ModPlatt: the highest ModPlatt score overall (i.e., the ModPlatt score of the highest-ranking candidate according to the ModPlatt score)
- “topsim”: For every instance of a dataset, the Tanimoto similarity of the highest-ranked candidate to the correct structure, based on the full (8925-bit) fingerprint of the molecule parsed from SMILES and the correct structure. For this analysis, SMILES that parse to the correct structure are *removed* from the result set. This permits to compare chemical accuracy of the model with, e.g, the training set, without biasing the analysis based on presence or absence of the correct structure in the dataset.
  ‐ ranked by raw score: the Tanimoto similarity of the highest-ranked candidate according to the model score, i.e. the probability under the model
  ‐ ranked by ModPlatt: the Tanimoto similarity of the highest-ranking candidate according to the ModPlatt score (*not* the highest Tanimoto similarity overall, as this cannot be known without knowing the correct structure and therefore cannot be used to rank a new prediction.)
- Chemical space coverage: As a reference, for each instance, all database search results with ModPlatt score equal to or higher than the correct structure were selected, using all instances for which the correct structure in the database was at rank 10 or better. For every instance, the *chemical space coverage* was calculated as the fraction of structures from the database reference which were in the set of predicted structures. This was used as a proxy measure to assess how well the chemical space produced by the model covers the chemical space in the database, while not being influenced by how well the predicted fingerprints match to the correct structure. The mean coverage per instance, the number of instances with 50% or more coverage (“% half”) and the number of instances with complete coverage (“% full”) were reported.

The following scores were calculated for an entire dataset:

- retrieval (“% found”): The fraction of instances for which the correct structure was present in the set of predicted structures
- top-*n* retrieval: The fraction of instances for which the correct structure was at rank *n* or better in the ordered set of predicted structures
  ‐ ranked by raw score: result set ordered according to the model score, i.e. the probability of the sequence under the model
  ‐ ranked by ModPlatt: result set ordered according to the ModPlatt score.

### De novo annotation of bryophyte metabolites

The dataset MTBLS709 was downloaded from the MetaboLights repository (ftp://ftp.ebi.ac.uk/pub/databases/metabolights/studies/public/MTBLS709). Using an R script, the 10436 MS^2^ spectra were consolidated by precursor (within 0.002 *m/z*) and similarity (> 0. 9) to a dataset of 6154 spectra. The dataset was submitted to GNPS for further clustering, molecular networking, initial annotation and visualization (https://gnps.ucsd.edu/ProteoSAFe/status.jsp?task=b8b481147b844ebda2481bf9656baec8). From the resulting clustered spectra set, the 576 spectra with *m/z* < 500 were selected. Spectra were processed with SIRIUS 4.4.29 for formula prediction (SIRIUS with profile Q-TOF, standard settings; ZODIAC with standard settings), fingerprint prediction and structure annotation (CSI:FingerID, search in “all databases except *in silico*”). The resulting dataset was filtered to retain only instances with high-confidence formula annotation ≥ 80% explained peaks, ≥ 90% explained intensity, ≥ 0. 9 ZODIAC score; 224 spectra) and used as input for *de novo* structure prediction. The structure candidates from database search and *de novo* prediction were both ranked by *ModPlatt* score and the top candidate selected. Instances where the top candidate from *de novo* prediction was a markedly better spectrum fit (*ModPlatt*_*MSNovelist*_ − *ModPlatt*_*database*_ > 50) were selected for further analysis (7 instances). The corresponding MS^2^ spectra were analyzed by hand, by library search (NIST MS version 2.4, in MS/MS Hybrid mode; product ion tolerance 0.02 *m/z*) with the NIST 20 library and the MassBank library, and using the NIST MS Interpreter (version 3.4.4; using protonated mass and 20 ppm).

We note that the implementation and parameters of our ModPlatt score are minimally different from the score implemented in SIRIUS 4.4.29, and in rare exceptions the top candidate found by our ModPlatt score might differ from the SIRIUS top candidate, however rescoring was necessary to achieve a comparison based on the same metric.

## Supplementary results

**Supplementary Figure 1:**
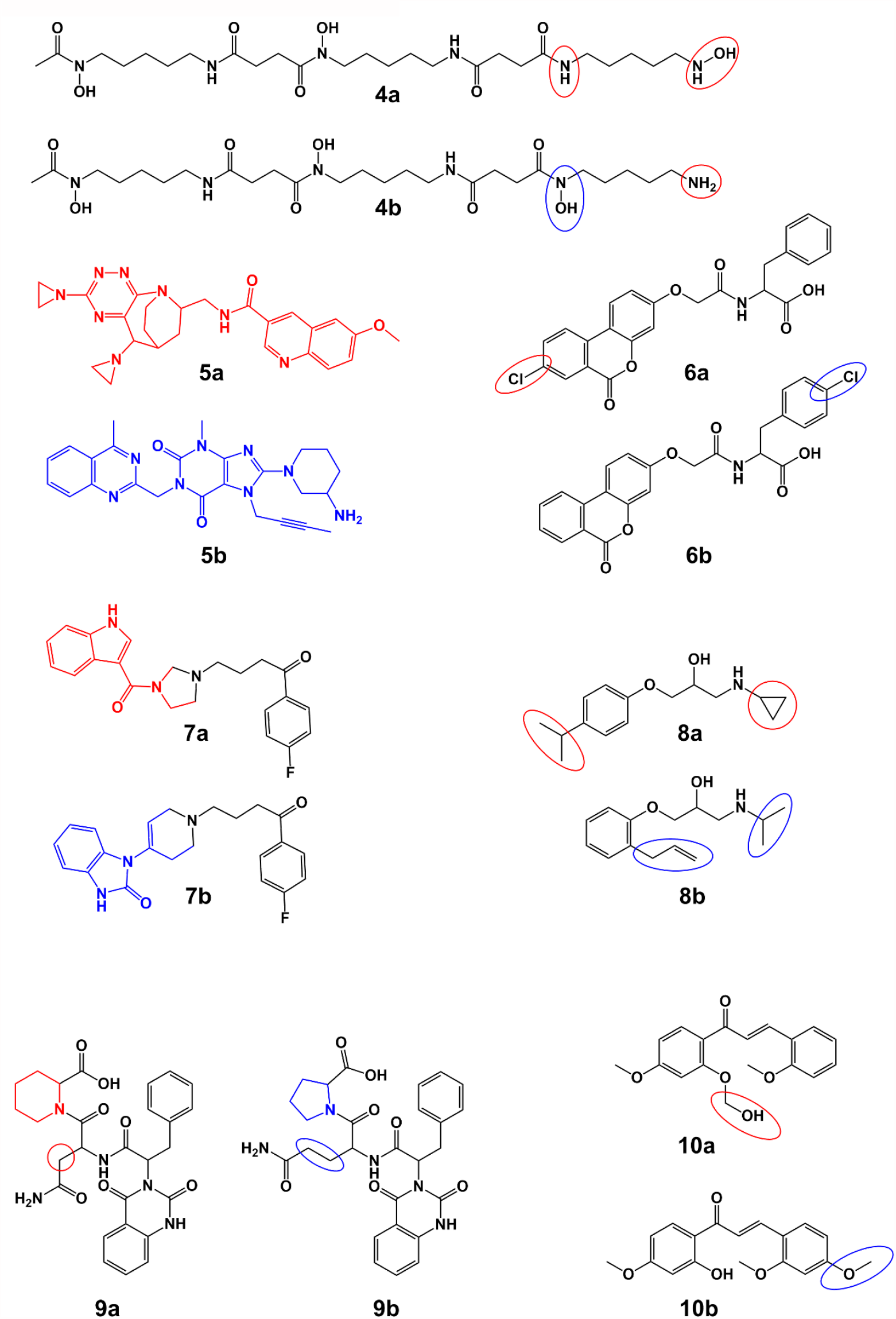
Seven randomly chosen incorrect MSNovelist predictions from the GNPS dataset. Structures 4-10a: de novo prediction; structures 4-10b: correct result. Red color marks sites predicted incorrectly by the model (or the entire molecule if the prediction was completely wrong), blue color marks the corresponding correct alternative.

**Supplementary Figure 2:**
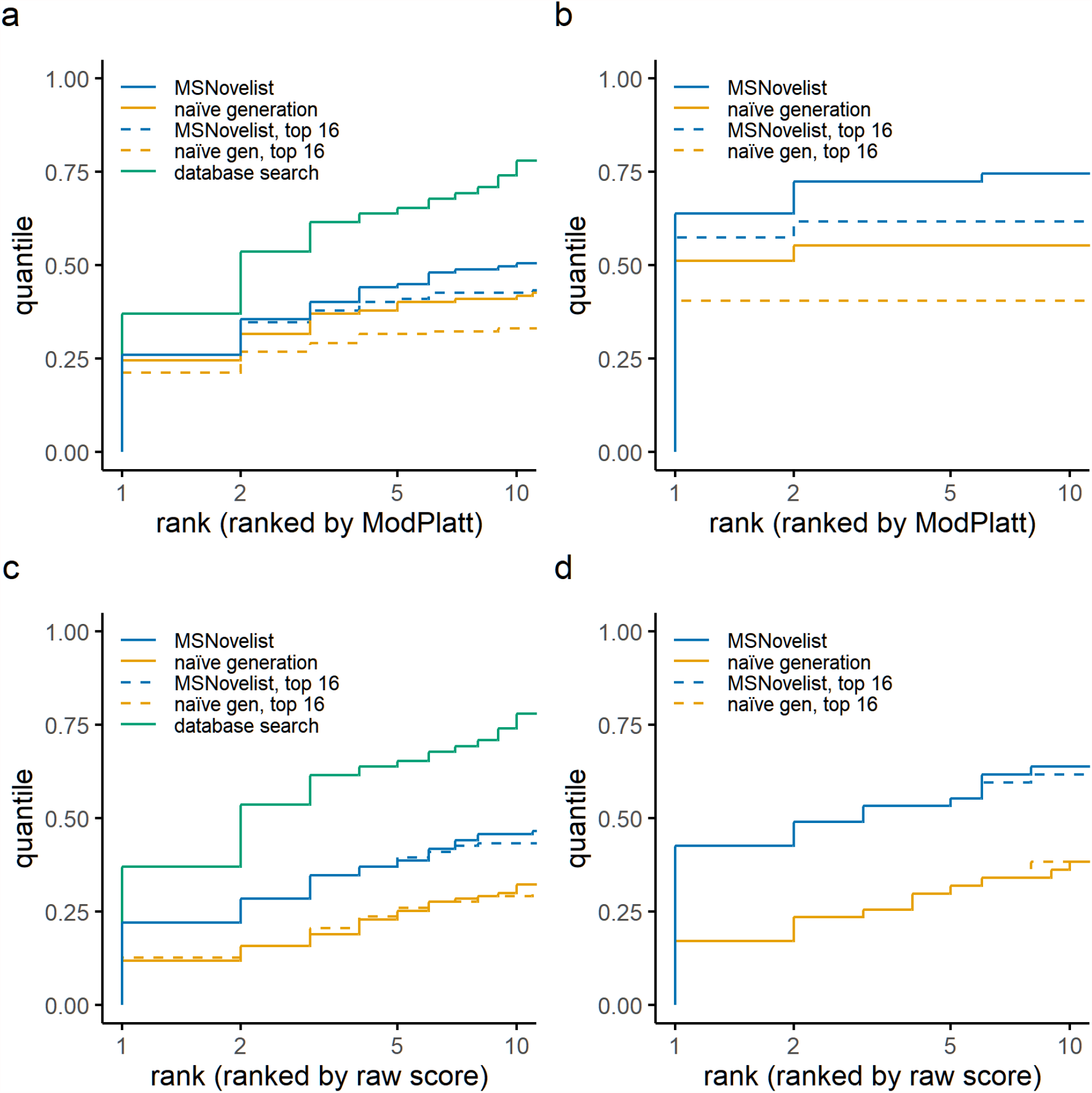
Model evaluation for CASMI dataset: Rank of correct structure in results for MSNovelist prediction (blue), and naïve generation (orange), generated with top-128 (solid) or top-16 beam search (dashed), with comparison to database search (SIRIUS 4.4.29 on PubChem; green). a), b): Candidates reranked by ModPlatt score; c), d): candidates ordered by raw score (model probability). a), c): CASMI dataset (n = 127). b), d): CASMI-top1 dataset (n = 43). Note that beam search with k = 16 is not identical to selecting the top-16 candidates from a beam search with k = 128, leading to small differences between non-reranked top-128 and top-16 results.

**Supplementary Figure 3:**
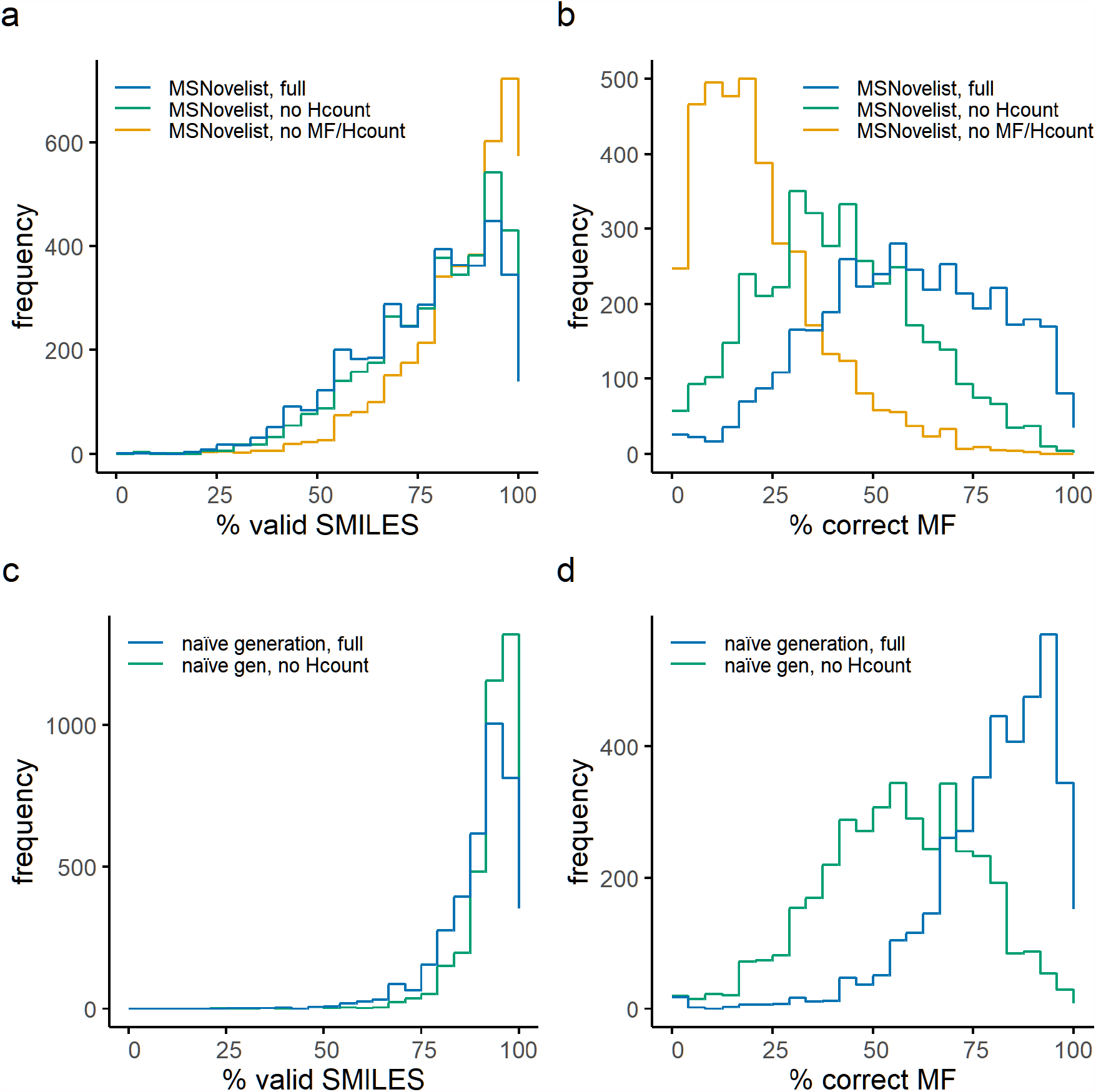
Validity of generated SMILES for MSNovelist model and naïve generation comparing full versus partial models, for GNPS dataset (n = 3863). a) Histogram of the fraction of valid SMILES per instance for full MSNovelist model (blue), MSNovelist model without hydrogen counting (green), and MSNovelist model without hydrogen counting and formula/grammar hinting (green). b) Histogram of the fraction of SMILES with correct molecular formula (versus all generated SMILES, including invalid SMILES) for full MSNovelist model, MSNovelist model without hydrogen counting, and MSNovelist model without hydrogen counting and formula/grammar hinting. c) Histogram of the fraction of valid SMILES per instance for full naïve model (blue) and naïve model without hydrogen counting (green). d) Histogram of the fraction of SMILES with correct molecular formula for full naïve model, and naïve model without hydrogen counting. Note: Naïve model with no hinting is not shown, since it generated ~ 100% valid SMILES but ~ 0% structures with correct formula, as expected (see Supplementary Table 1).

**Supplementary Figure 4:**
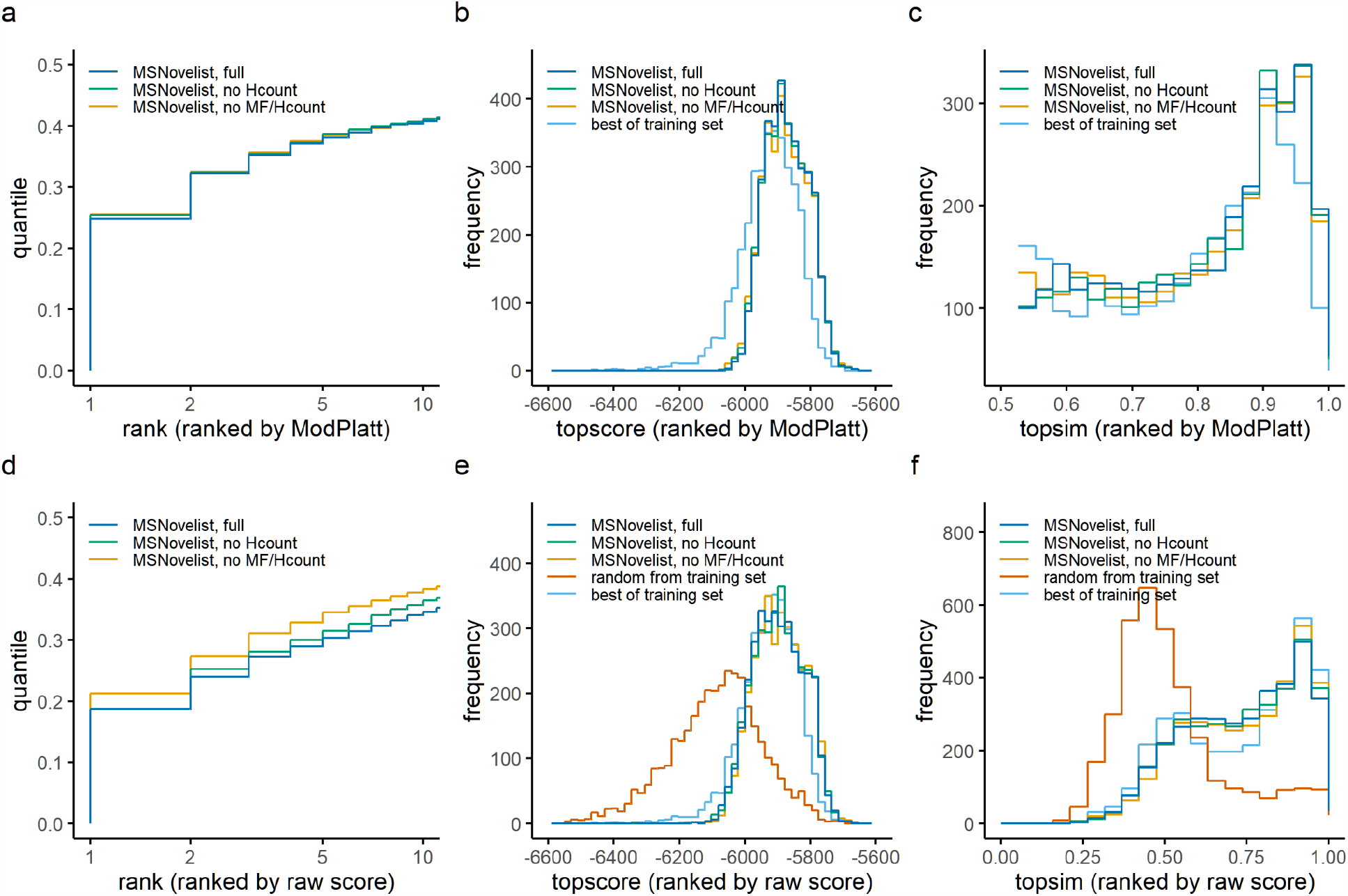
Evaluation of de novo model with and without formula hinting and hydrogen count, for GNPS dataset (n = 3863). a) Rank of correct structure in results for full de novo model (blue), de novo model without hydrogen counting (green), and de novo model without hydrogen counting and formula/grammar hinting (orange). b) ModPlatt score of top candidate (topscore) ranked by ModPlatt score, for full MSNovelist model, MSNovelist model without hydrogen counting, and MSNovelist without hydrogen counting and formula/grammar hinting, versus best candidate in training set (light blue). c) Tanimoto similarity of best incorrect candidate to correct structure (topsim) for full MSNovelist model, MSNovelist model without hydrogen counting, and MSNovelist model without hydrogen counting and formula/grammar hinting, versus best candidate in training set. d), e), f): idem, ordered by raw score (model probability); versus random choice from training set (red).

**Supplementary Figure 5:**
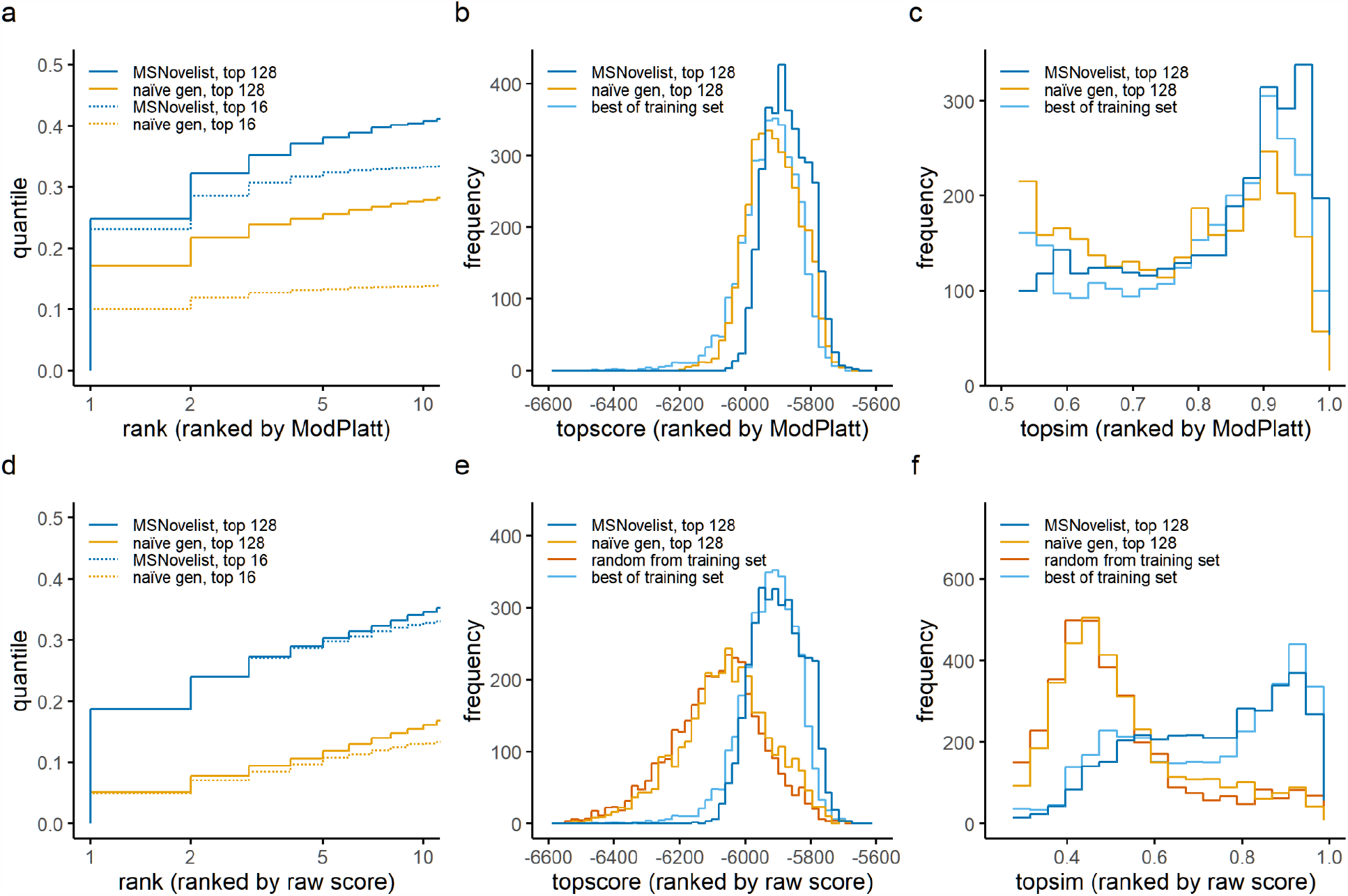
Evaluation of de novo generation versus naïve generation for top-128 and top-16 beam search, for GNPS dataset (n = 3863). a) Rank of correct structure in results for de novo prediction (blue), and naïve generation (orange), generated with k = 128 (solid) or k = 16 (dotted). b) ModPlatt score of top candidate (topscore) ranked by ModPlatt score, for MSNovelist prediction and naïve generation, with k = 128 or k = 16, versus best candidate from training set (light blue). c) Tanimoto similarity of best incorrect candidate to correct structure (topsim) for MSNovelist prediction and naïve generation, with k = 128 or k = 16, versus best candidate from training set. d), e), f): idem, ordered by raw score (model probability); versus random choice from training set (red).

**Supplementary Figure 6:**
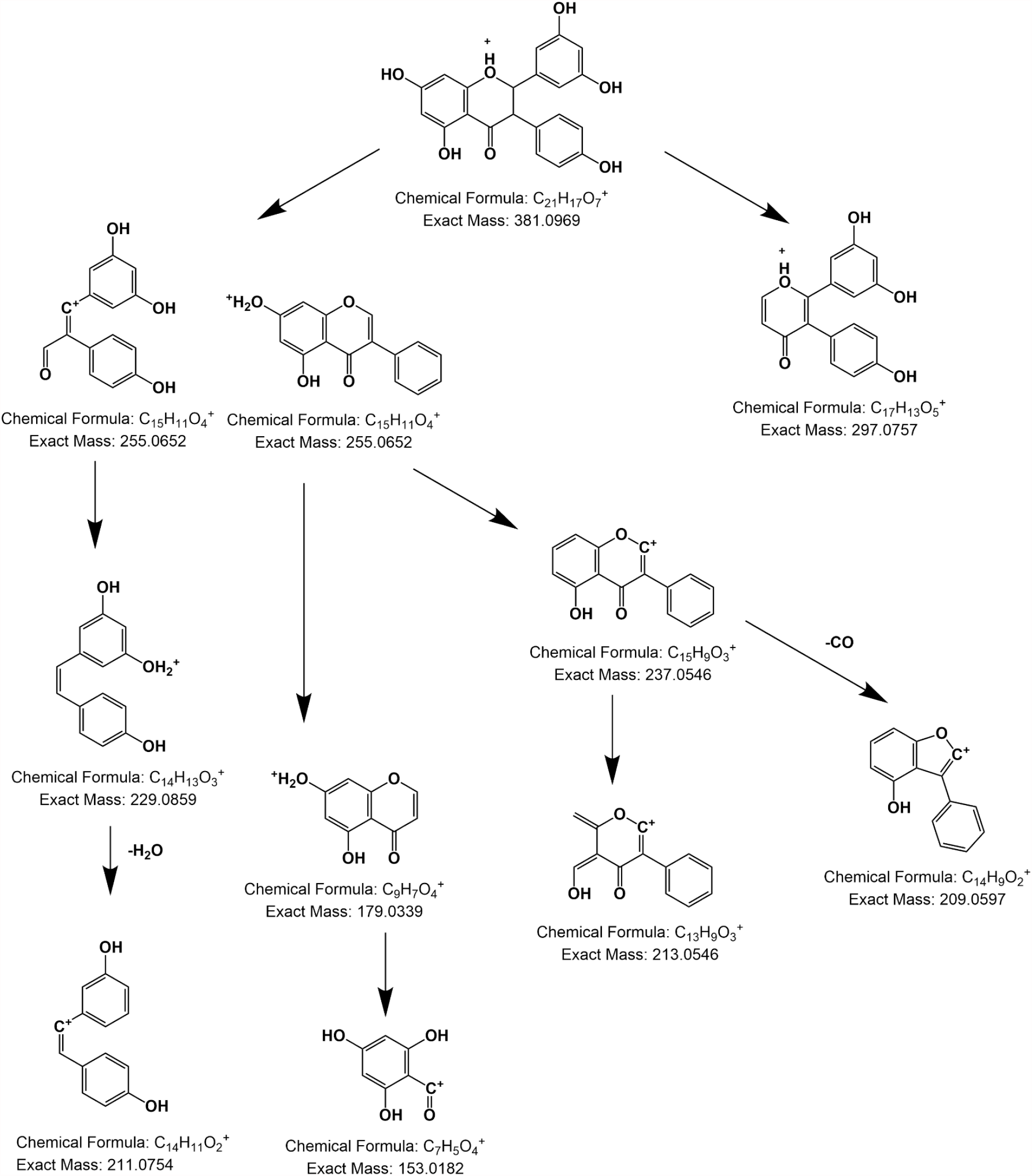
Proposed fragmentation for structure 377a.

Supplementary Tables 1-4: see “SuppTables1-4.xlsx”

Supplementary Tables 5-13: see “SuppTables5-13.xlsx”

